# Longitudinal trajectories of the neural encoding mechanisms of speech-sound features during the first year of life

**DOI:** 10.1101/2023.11.24.567803

**Authors:** Marta Puertollano, Teresa Ribas-Prats, Natàlia Gorina-Careta, Siham Ijjou-Kadiri, Sonia Arenillas-Alcón, Alejandro Mondéjar-Segovia, María Dolores Gómez-Roig, Carles Escera

## Abstract

Infants quickly recognize the sounds of their mother language, perceiving the spectrotemporal acoustic features of speech. However, the underlying neural machinery remains unclear. We used an auditory evoked potential termed frequency-following response (FFR) to unravel the neural encoding maturation for two speech sound characteristics: voice pitch and temporal fine structure. The FFR was elicited by a two-vowel stimulus (/oa/) in a sample of 41 healthy-term neonates, tested at birth and retested at the ages of six and twelve months. Results revealed a reduction in neural phase-locking time to the stimulus envelope from birth to six months, remaining stable thereafter. While neural encoding of voice pitch remained consistent, temporal fine structure encoding matured rapidly from birth to six months, without further improvement from this age onwards. Results highlight the critical importance of the first six months in the maturation of neural encoding mechanisms, crucial for phoneme discrimination during early language acquisition.

## 1. Introduction

Infants show a native talent for language acquisition even since the very early stages of development. Behavioral evidence has shown that infants follow similar developmental trajectories in the acquisition of their mother tongue regardless of their culture and language, evolving from babbling (5 to 10 months; Kuhl, 2004) to their first words utterances by the age of 10 to 18 months (Feldman, 2019). Yet, there is a lack of consensus regarding the neural mechanisms supporting this talent. Starting from the tuning to the phonetic repertoire of the mother tongue, language acquisition entails a sophisticated fine-grained neural machinery across the entire auditory system to capture the complex spectro-temporal acoustic features that characterize the speech sounds.

Despite postnatal hearing experience is essential for an adequate auditory and language development, neonates are born with a wide range of universal speech perception abilities that allow them to acquire any language. For instance, neonates can discriminate different languages they have not been exposed to if these are rhythmically different (Byers-Heinlein et al., 2010; Mehler et al., 1988; Nazzi et al., 1998). They can also encode the pitch of a speech sound in an adult-like manner (Arenillas-Alcón et al., 2021; Jeng et al., 2011, 2016), as well as recognize their mothers’ voice (Decasper & Fifer, 1980; Hepper et al., 1993) and melodies they have been exposed to during pregnancy (Granier-Deferre et al., 2011).

Yet to imitate speech and acquire a mother tongue, non-identical sounds must be perceived as falling into either separate or equivalent phonetic categories in a language-dependent manner. This competence emerges a few months after birth, in parallel with the myelination trajectory of the auditory pathway (Moore et al., 1995) and the exposure to their specific native language (Kuhl et al., 1992, 2003; Rivera-Gaxiola et al., 2005), and it is mediated by statistical learning (Jusczyk et al., 1994). Previous studies using behavioral paradigms demonstrated that by the age of six months, babies are able to perceive the variability inherent in each phonetic unit. This ability enables them to identify vowels typical of their mother tongue, which alters phonetic perception towards a native-like model (Kuhl et al., 1992; Maye et al., 2002). Vowel discrimination at this stage, as measured by the conditioned head turn paradigm, may serve as a predictor of future infant language abilities at the age of 13, 16 and 24 months (Tsao et al., 2004). A decline in the discrimination ability for nonnative contrasts in language becomes evident by the end of the first year of life, as assessed through behavioral paradigms (Werker and Tees, 1984) and event-related potentials (Rivera-Gaxiola et al., 2005; Tsao et al., 2006). It is at this moment that infants start to develop an adult-like attunement to their native language phoneme repertoire (Cheour et al., 1998; Kuhl et al., 2006; Werker et al., 1981).

Research on neural mechanisms underlying the acquisition of speech sounds has benefited from advances in the use of neonatal and infant brain potentials (Hervé et al., 2022). One such evoked potential is the mismatch negativity (MMN), generated by acoustic and linguistic changes (Kujala et al., 2023). Another recently growing body of research is exploring the so-called frequency-following response (FFR). The FFR is a non-invasive auditory evoked potential that is elicited by periodic complex stimuli such as speech or music, and reflects neural activity from the auditory nerve to the cortex (Coffey et al., 2019; Gorina-Careta et al., 2021). It mimics the eliciting stimulus, thus providing a unique snapshot into the neural encoding of the two distinctive features that characterize the speech sounds: voice pitch, associated with its fundamental frequency (F_0_), and the temporal fine structure, associated with its formants (Aiken & Picton, 2008; Krizman & Kraus, 2019). The FFR has been studied in infancy to characterize normal and abnormal developmental trajectories of neural speech encoding (Banai et al., 2005, 2009; Cunningham et al., 2001), as these appear disrupted in children with literacy impairments, including dyslexia (Hornickel & Kraus, 2013), specific language impairment (Basu et al., 2010), and autism (Font-Alaminos et al., 2020; Otto-Meyer et al., 2018; Russo et al., 2008).

The FFR has also been explored during the first months of life in several cross-sectional studies with different age periods of interest, as an attempt to describe its typical trajectory along early development. A decrease in neural conduction times and neural phase-locking onset has been already observed at the early age of 45 days (Ferreira et al., 2021), with further shortenings until the age of ten months (Anderson et al., 2015). An adult-like voice pitch encoding at birth has also been reported (Arenillas-Alcón et al., 2021; Jeng et al., 2016), with a more robust neural representation with age across the first year of life (Jeng et al., 2010; Ribas-Prats et al., 2023b; Van Dyke et al., 2017). The maturation of neural encoding of temporal fine structure components, as assessed through neural responses to the high-frequency formants and harmonics, begins as early as the first month of life (Ribas-Prats et al., 2023b) and continuous to develop until the age of ten months (Anderson et al., 2015).

However, the studies reviewed above provide an incomplete view of the developmental trajectory of speech-sounds neural encoding mechanisms during the first year of age. Behavioral paradigms impose constraints on disentangling the neural underpinnings of speech perception. Furthermore, cross-sectional designs adopted by previous electrophysiological studies offer a limited approach to characterize the neural correlates of speech development. The present longitudinal study was set to provide a pioneering and comprehensive picture of the maturational pattern of the neural mechanisms involved in encoding two distinct speech-sound features during the first postnatal year, as reflected in the FFR: voice pitch, as represented by its fundamental frequency, and speech temporal fine structure, corresponding to its formants. We hypothesized an enhancement in the neural encoding of these two speech-sound features as a function of age, starting from birth to six months and further continuing from six months to the age of one year. Neural phase-locking onset was also expected to decrease due to the well-known myelination process of the auditory pathway during the first year of life.

## 2. Methods

### 2.1 Participants

Sixty-six healthy-term neonates (31 females; mean gestational age at birth = 39.72 ± 0.95 weeks; mean birth weight = 3295.45 ± 308.14 grams; mean age at evaluation = 1.94 ± 1.73 days after birth) were recruited at the Sant Joan de Déu Barcelona Children’s Hospital (Catalonia, Spain). Fifty-four of them (27 females; mean gestational age at birth = 39.73 ± 0.97 weeks; mean birth weight = 3309.17 ± 313.08 grams; mean age at evaluation = 1.81 ± 1.28 days after birth) were followed-up at the age of six months (aged 5.53 to 7.77 months after birth; mean = 6.42 ± 0.43 months). Forty-one neonates (21 females; mean gestational age at birth = 39.71 ± 0.91 weeks; mean birth weight = 3293.29 ± 310.27 grams; mean age at evaluation = 1.75 ± 1.09 days after birth) were further followed-up at the age of one year, and hence completed all stages of the study: at the age of six months (aged 5.53 to 7.5 months after birth; mean = 6.41 ± 0.37 months) and at twelve months of age (aged 11.97 to 13.7 months after birth; mean = 12.60 ± 0.40 months).

All neonates were born after low-risk gestations, without either pathologies or risk factors for hearing impairment, following the Joint Committee on Infant Hearing guidelines (2019). Apgar scores were higher than 7 at 1 and 5 minutes after birth and, in all cases, birth weight was adequate for their gestational age (Figueras & Gratacós, 2014). Furthermore, all infants had passed the universal hearing screening test as part of the standard medical routine, based on an automated auditory brainstem response system to ensure auditory pathway health (ALGO 3i, Natus Medical Incorporated, San Carlos, CA). To confirm the integrity of the auditory pathway, an auditory brainstem response (ABR) to a click stimulus (10 µs; delivered monaurally to the right ear at 60 dB SPL at a rate of 19.30 Hz, for a total of 4000 averaged sweeps) was also obtained from every neonate.

The study was approved by the Bioethics Committee of SJD Barcelona Children’s Hospital (Internal review board ID: PIC-185-19). A written informed consent was obtained from all parents or legal guardians prior to the data collection in accordance with the Code of Ethics of the World Medical Association (Declaration of Helsinki). The data that supports the findings of this study and the code used for data analysis are available upon request to the corresponding author.

### 2.2 Stimulus

To obtain the FFR, a two-vowel /oa/ stimulus was used (see Fig. 1A), as previously designed in our laboratory by Arenillas-Alcón et al. (2021). The synthesized stimulus had a total duration of 250 ms, with a steady F_0_ at 113 Hz for its first 160 ms and a linearly rising F_0_ from 113 to 154 Hz during its last 90 ms (from 160 to 250 ms). The specific first formants (F_1_) of the vowels (/o/ F_1_ = 452 Hz; /a/ F_1_ = 678 Hz) were chosen for the stimulus design as those are found in the prototypical phonetic repertoires of both Catalan and Spanish languages (Alarcos Llorach, 1965; Martí i Roca, 1986). The stimulus was presented monaurally to the right ear at a rate of 3.39 Hz and an intensity of 60 dB SPL in alternating polarities through Etymotic shielded earphones of 300 Ω (ER, Elk Grove Village, IL, EEUU) connected to a Flexicoupler® disposable adaptor (Natus Medical Incorporated, San Carlos, CA).

**Fig. 1.**
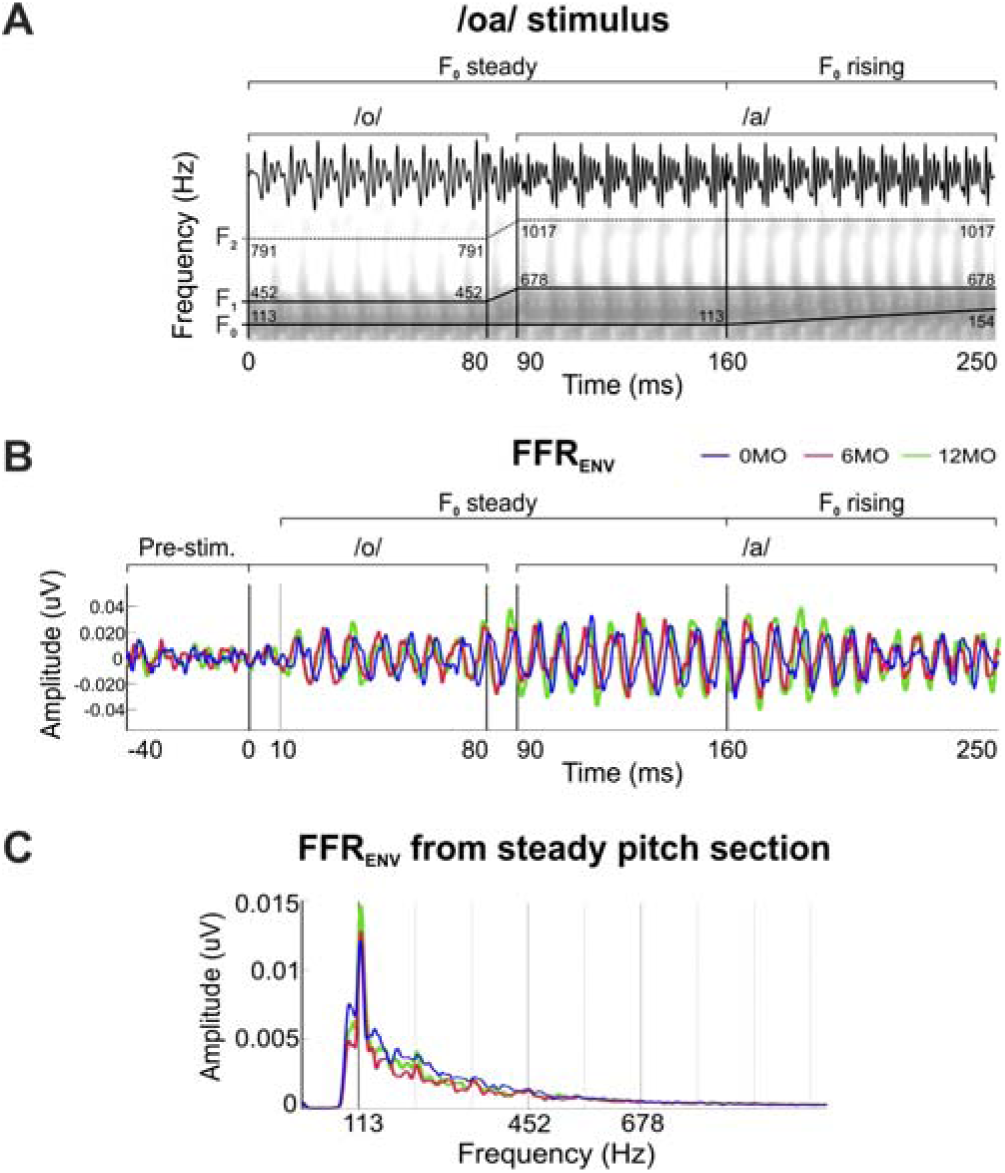
(**A**) Temporal and spectral plots of the /oa/ stimulus with a schematic representation of its formant structure. F_0_, F_1_ and F_2_ are defined for the different stimulus sections. (**B**) Grand-averaged FFR_ENV_ waveforms in the time domain from the 41 infants recorded at birth (0MO, blue), six months (6MO, red) and twelve months of age (12MO, green), obtained after averaged neural response polarities. (**C**) Amplitude FFR_ENV_ spectra extracted from the analyzed steady part of the stimulus (10-160 ms).

### 2.3 Procedure

After a successful universal hearing screening test, the ABR and subsequently the FFR were recorded at the hospital room while the newborns were sleeping in their crib, following the same protocol used in previous studies (see Arenillas-Alcón et al., 2021, 2023; Ribas-Prats et al., 2019, 2022, 2023a, 2023b; for a review see Gorina-Careta el al., 2022). Recording was interrupted to any sign of discomfort or sleep disruption and it was resumed when the newborn was asleep again. Infants that successfully completed the neonatal FFR session were invited to two successive follow-up sessions: at six and twelve months of age. Out of the 66 recruited neonates only 41 completed the two follow-up sessions, yielding a failure to complete the longitudinal study at one year in 39.7% of the participants. The infants that returned were retested at six and at twelve months of age at a hospital dispensary, keeping the baby either asleep or as calm as possible during the recording in order to ensure the highest quality of the EEG data. The total mean duration of the sessions was around 30 minutes, including a preparation time of around 5 minutes, 20 minutes of recording (four /oa/ stimulus blocks × 1000 sweeps × 295 ms SOA), and the additional time for the rejected sweeps.

### 2.4 Data acquisition

ABR and FFR recordings were carried out using a SmartEP platform connected to a Duet amplifier, including the cABR and Advanced Hearing Research modules (Intelligent Hearing Systems, Miami, Fl, USA). Three disposable Ag/AgCl electrodes located in a vertical montage were employed for the recording, with the active electrode placed at Fpz, ground at the forehead and reference at the right mastoid. Impedances were kept below 10 kΩ for all electrodes. The continuous FFR signal was acquired at a sampling rate of 13333 Hz with an online bandpass filter between 30 and 1500 Hz. Online data was epoched from -40.95 (for the baseline period) to 249.975 ms (relative to the stimulus onset). An automated online rejection of artifacts was applied, excluding any sweep with voltage values exceeding ± 30 µV. The percentage of rejected sweeps was kept below 7% at birth, 33% at six months and 39% at twelve months.

### 2.5 Data processing

An offline bandpass filter from 80 to 1500 Hz was applied for FFR analysis. Neural responses to alternating polarities were averaged [(Condensation + Rarefaction)/2] to emphasize the FFR components related to the stimulus envelope (FFR_ENV_) and to minimize the involvement of putative cochlear microphonics. In addition, to assess the neural encoding of the vowels’ F_1_ and minimizing the contribution of envelope related activity (Aiken & Picton, 2008; Krizman & Kraus, 2019), the FFR temporal fine structure (FFR_TFS_) was analyzed by subtracting the neural response to the alternating polarities [(Rarefaction–Condensation)/2]. Only the steady part of the stimulus was considered for the analysis. Thus, the FFR_TFS_ (spectral peaks corresponding to F_1_) was analyzed separately for the /o/ section (10 to 80 ms, F_0_ = 113 Hz, F_1_ = 452 Hz) and for the /a/ steady section (90 to 160 ms, F_0_ = 113 Hz, F_1_ = 678 Hz).

FFR parameters were evaluated using custom scripts from Matlab R2019b (The Mathworks Inc., 2019) used in previous studies performed in our laboratory (Arenillas-Alcón et al., 2021; Ribas-Prats et al., 2019). A comprehensive description is given below for the three parameters extracted and tested separately for the different frequencies of interest (for a detailed description, see Arenillas-Alcón et al., 2021; Ribas-Prats et al., 2019).

#### Neural lag

In order to assess the neural transmission delay occurring along the auditory pathway, the neural lag value was obtained. This parameter accounts for the time lag between the stimulus presentation and the neural phase-locking onset (Arenillas-Alcón et al., 2021; Jeng et al., 2010; Ribas-Prats et al., 2019). It was computed from the cross-correlation between the stimulus and the neural response as the time shift that corresponded to the maximum cross-correlation magnitude.

#### Spectral amplitude

In order to analyze the neural-phase locking magnitude at the frequency of interest (F_0_, 113 Hz; /o/ F_1_, 452 Hz; /a/ F_1_, 678 Hz), spectral amplitude was obtained as an indicator of the response strength at that given frequency (Arenillas-Alcón et al., 2021; Ribas-Prats et al., 2019; White-Schwoch et al., 2015b). To obtain the FFR frequency decomposition, spectral amplitude was calculated after applying the Fast Fourier Transform (FFT; Cooley & Tukey, 1965), by computing the mean amplitude within a ±5 Hz frequency window centered at the frequency peak of interest. Spectral amplitude at F_0_ was retrieved from the FFR_ENV_ corresponding to the /oa/ steady section (10 to 160 ms) to assess voice pitch encoding of the speech-sound stimulus. Spectral amplitudes at the stimulus F_1_ frequencies were extracted separately from the FFR_TFS_ corresponding to the /o/ section (10 to 80 ms) and the /a/ steady section (90 to 160 ms).

#### Signal-to-noise ratio

Signal-to-noise ratio (SNR) at the frequency peak of interest was calculated in order to estimate the FFR relative spectral magnitude. It was computed by dividing the spectral amplitude value obtained for the given frequency of interest (±5 Hz window centered at the peak of interest) by the mean amplitude of its two flanks (28 Hz windows centered at ±19 Hz from the frequency of interest). Normalized values were calculated for SNR according to the formula (see Arenillas-Alcón et al., 2021):

> SNR = 10 ∗ log10 (Signal spectral power ⁄ Noise spectral power).

SNR at F_0_ was extracted from the FFR_ENV_ to evaluate voice pitch encoding. SNRs at vowels F_1_ were retrieved from the FFR_TFS_ to assess the formant structure encoding of the auditory stimulus and analyzed following the same procedure as for the spectral amplitude parameter (i.e., the values were extracted separately from the neural responses to the vowel sections).

### 2.6 Statistical analysis

Statistical analyses were performed using SPSS 25.0 (IBM Corp, Armonk, NY). Descriptive statistics are presented for each parameter as median and interquartile range for each time of measurement (see Table 1). Results were considered statistically significant when p < .05. Normality was assessed with Shapiro-Wilk’s test and, as all parameters followed a non-normal distribution, Friedman’s test was applied. After a given significant result, Wilcoxon signed-rank test was employed to explore each time point measurements pair.

**Table 1.** Descriptive statistics expressed as median (IQR, interquartile range), and Friedman test comparison between the 41 neonates recorded at birth (0-MO) and their retest at the age of six (6-MO) and twelve months (12-MO) for each FFR parameter assessed. Wave V amplitude and latency values at birth are also depicted for the extended 66 neonatal sample as mean (SD).

| Measures | 0-MO | 6-MO | 12-MO | Friedman test | df | *p* value |
| --- | --- | --- | --- | --- | --- | --- |
| **Wave V (N = 66)** |  |  |  |  |  |  |
| Amplitude (μV) | 0.08 (0.14) | - | - | - | - | - |
| Latency (ms) | 8.50 (0.38) | - | - | - | - | - |
| **FFR (N = 41)** |  |  |  |  |  |  |
| Neural lag (ms) | 8.03 (2.03) | 6.68 (1.95) | 6.53 (1.76) | 22.84 | 40 | <.001 |
| **FFR<sub>ENV</sub>** |  |  |  |  |  |  |
| Spectral amplitude F<sub>0</sub> (μV) | .008 (.008) | .009 (.012) | .013 (.014) | 2.10 | - | .35 |
| SNR F<sub>0</sub> (dB) | 3.99 (5.69) | 4.75 (7.90) | 6.41 (5.41) | 1.81 | - | .41 |
| **FFR<sub>TFS</sub>** |  |  |  |  |  |  |
| Spectral amplitude at /o/ F<sub>1</sub> (10-80 ms; μV) | .0018 (.002) | .0031 (.005) | .0027 (.005) | 4.44 | - | .10 |
| SNR at /o/ F<sub>1</sub> (10-90ms; dB) | 1.38 (4.88) | 4.83 (6.98) | 3.52 (4.96) | 15.85 | - | <.001 |
| Spectral amplitude at /a/ F<sub>1</sub> (90-160 ms; μV) | .0008 (.001) | .0012 (.002) | .0017 (.002) | 6.78 | - | .034 |
| SNR at /a/ F<sub>1</sub> (90-160 ms; dB) | 1.73 (4.08) | 2.61 (5.99) | 3.81 (3.79) | 7.37 | - | .025 |

In addition, to ensure that the spectral amplitude and SNR measurements obtained for the stimulus F_1_ where specific to the corresponding stimulus vowel section (i.e., 452 Hz for the /o/ vowel, and 678 Hz for the /a/ vowel), as well as its possible interaction with age, a repeated measures Analysis of Variance (rmANOVA) test was performed. For that, the variables Age (0, 6 and 12 months) and Stimulus Section (/o/ and /a/) were chosen as within-subject factors. Bonferroni correction was applied to adjust *p*-values for multiple pairwise comparisons. Greenhouse-Geisser correction was used when the assumption of sphericity was violated. Partial eta squared (ηp2) was reported as a measure of effect size.

An ABR to a click stimulus was obtained from every neonate before the FFR recording to confirm the integrity of the auditory pathway. All recruited infants (N = 66) had an identifiable wave V peak at birth, with a mean latency of 8.50 (± 0.38 SD) ms and a mean amplitude of 0.08 (± 0.14 SD) μV (Table 1). Values were similar to those previously reported at the literature (Arenillas et al., 2021; Ribas-Prats et al., 2019; Stuart et al., 1994).

In order to unravel the maturational pattern of the neural encoding of speech sounds features during the first year of life, FFRs elicited by the /oa/ stimulus were collected from the sample of forty-one neonates that completed the follow-up at the ages of six and twelve months. The corresponding grand-average FFR_ENV_ and FFR_TFS_ waveforms are shown in Fig. 1B and Fig. 2A respectively. Table 1 depicts the descriptive statistics and results from the Friedman test comparison for all FFR parameters evaluated at the three developmental stages (i.e., 0, 6 and 12 months).

**Fig. 2.**
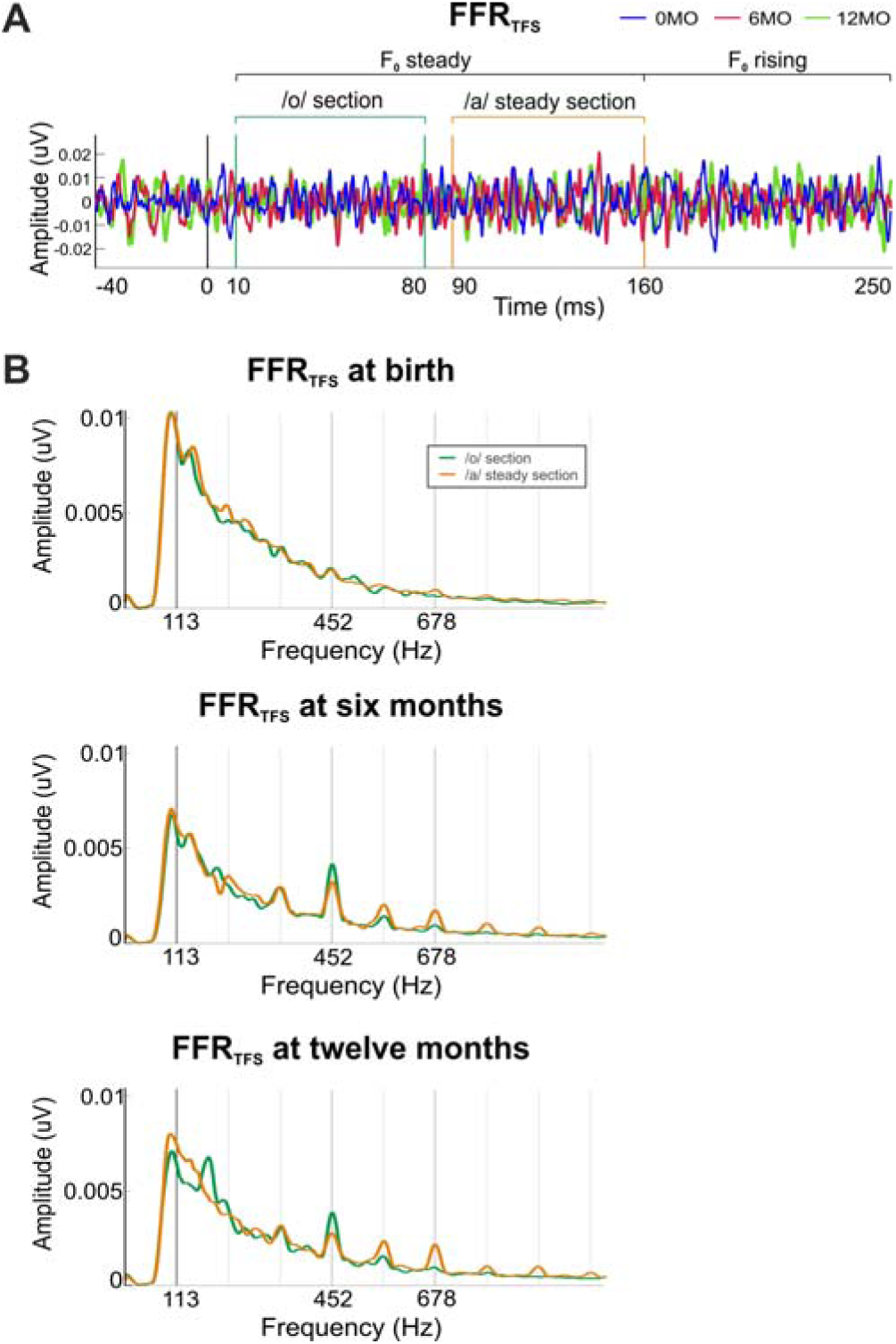
(**A**) Time-domain grand-averaged FFR_TFS_ waveforms extracted after substracting neural responses to alternating stimulus polarities from the 41 infants recorded at birth (0MO, blue) and retested at six months (6MO, red) and twelve months of age (12MO, green). (**B**) Amplitude FFR_TFS_ spectra obtained for the two vowel sections: /o/ (green), /a/ (orange).

### 3.1 Neural lag

Results revealed a consistently shortened neural phase-locking onset as a function of age (X^2^(2) = 27.84, *p* < .001; see Fig. 3A). Post-hoc analyses revealed shorter neural lag at both six (*Mdn* = 6.68; *z* = -3.51, *p* < .001) and twelve months of age (*Mdn* = 6.53; *z* = -4.39, *p* < .001) in comparison to that at birth (*Mdn* = 8.03). Neural transmission delay at the age of six and twelve months were similar (*z* = -.97, *p* = .33). To further support the results obtained, statistical analyses were repeated for the entire sample that completed the follow-up session at six months of age (N = 54). Similar results were obtained for neural transmission delay (i.e., shortened neural lag at six months; *z* = - 3.79, *p* = < .001; see Table 2).

**Fig. 3.**
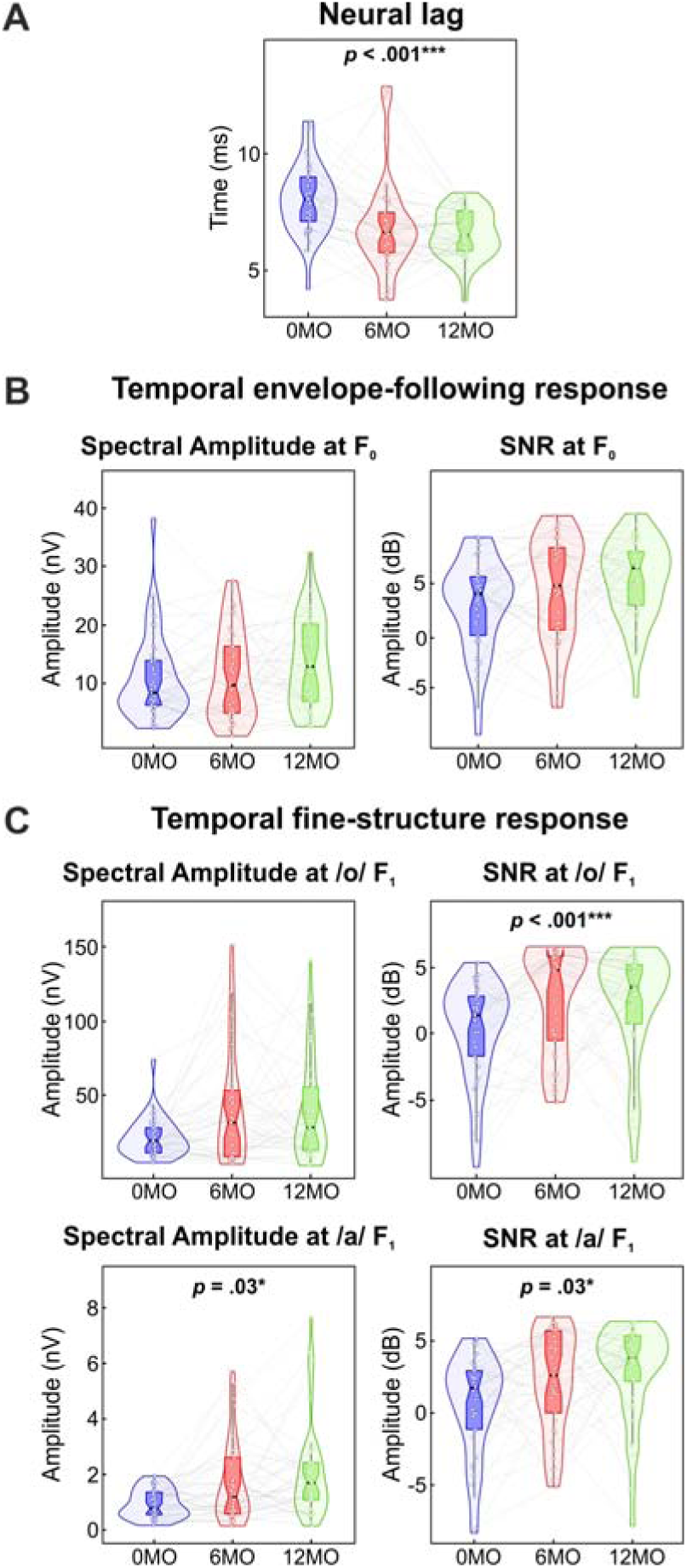
Data distribution from the 41 infants recorded at birth (0MO, blue) and retested at six (6MO, red) and twelve months of age (12MO, green). Violin plots are depicted for (**A**) the neural lag, (**B**) the FFR_ENV_ obatined to the steady part of the stimulus (10-160 ms), and (**C**) the FFR_TFS_ corresponding to /o/ (upper panel) and /a/ (lower panel) stimulus sections.

**Table 2.**
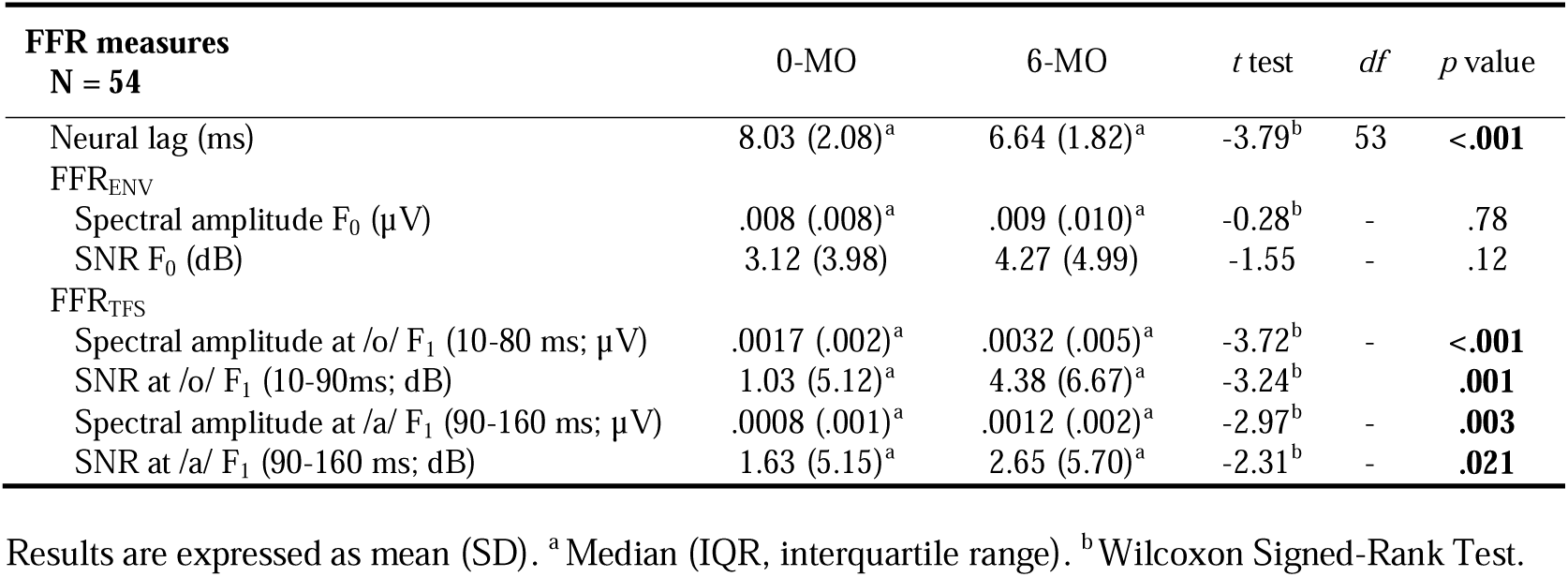
Descriptive statistics and comparison between the sample of 54 neonates recorded at birth (0-MO) and their retest at the age of six (6-MO) for each FFR parameter assessed.

### 3.2 Temporal envelope-following response

Grand-average FFR_ENV_ waveforms were obtained at each developmental stage (see Fig. 1B). The corresponding frequency spectrum for averaged polarities is shown in Fig. 1C. The strength of the stimulus F_0_ neural representation along the first year of life was assessed by means of the spectral amplitude and SNR parameters computed on the FFR_ENV_. Statistical analyses revealed no significant differences across age (at birth, at six and at twelve months) in neither spectral amplitude parameter (X^2^(2) = 2.10, *p* = .35) or in SNR (X^2^(2) = 1.81, *p* = .41; see Fig. 3B). Results remained statistically similar for both parameters in the analysis with the extended fifty-four infants sample that could complete the recording at the age of six months (i.e., spectral amplitude, *z* = -.28, *p* = .78; SNR, *t_(53)_* = -1.55, *p* = .13).

### 3.3 Temporal fine-structure response

The maturation of neural mechanisms for the encoding of the speech-sound formant structure along the first year of life was analyzed from the FFR_TFS_. Grand-average FFR_TFS_ waveforms are illustrated in Fig. 2A for each developmental stage. In order to evaluate phase-locking at the stimulus F_1_, neural responses to each stimulus vowel section were assessed separately. Fig. 2B illustrates the frequency spectrum corresponding to both vowel sections at birth, six and twelve months of age. Spectral amplitudes and SNRs were retrieved selecting the spectral peaks corresponding to the frequency of interest according to each vowel (452 Hz for the /o/; 678 Hz for the /a/). Fig. 3C depicts spectral amplitude and SNR values for the FFR_TFS_ along the three developmental stages.

#### Neural encoding of the /o/ vowel F_1_

Spectral amplitude and SNR at /o/ vowel F_1_ (452 Hz) were analyzed at the corresponding stimulus /o/ vowel section. No differences were observed for spectral amplitude between the three stages of development (X^2^(2) = 4.44, *p* = .1; see Table 1). For the SNR, significant differences were obtained (X^2^(2) = 15.85, *p* < .001), with larger values at six months (*Mdn* = 4.83; *z* = -2.73, *p* = .006) and twelve months of age (*Mdn* = 3.52; *z* = -2.60, *p* = .009) in comparison to birth (*Mdn* = 1.38); SNR values at six and twelve months of age were similar (*z* = -.72, *p* = .47). Wilcoxon signed-rank test assessed for the extended six-months old sample (i.e., fifty-four infants) revealed larger values at six months of age in the two parameters assessed (spectral amplitude, *z* = -3.72, *p* < .001; SNR, *z* = -3.24, *p* = .001; see Table 2).

To investigate the specificity of the neural encoding of the formant structure corresponding to each of the two vowels of the /oa/ stimulus, and its possible interaction with age, a two-way rmANOVA test was conducted with the factors Age (0, 6 and 12 months) and Stimulus Section (/o/ and /a/) on the spectral amplitude and its SNR at 452 Hz, corresponding to the /o/ F_1_ (see Fig. 4). Spectral amplitude results revealed a main effect of stimulus section (*F*_(1,40)_ = 7.96, *p* = .007, ηp2 = .17), with higher spectral amplitudes for the /o/ section (*M* = .0033 ± < .001) compared to the /a/ section (*M* = .0026 ± < .001). A main effect of age was also observed (*F*_(2,80)_ = 4.50, *p* = .014, ηp2 = .10), with larger amplitudes at the age of six months (*M* = .004 ± .001; *p* = .012) and the age of twelve months (*M* = .003 ± < .001; *p* = .048) in comparison to birth (*M* = .002 ± < .001). No significant developmental changes between six and twelve months-old stages were found (*p* = 1). Interaction between age and stimulus section was not significant (*F*_(2,80)_ = 2.95, *p* = .06, ηp2 = .07).

**Fig. 4.**
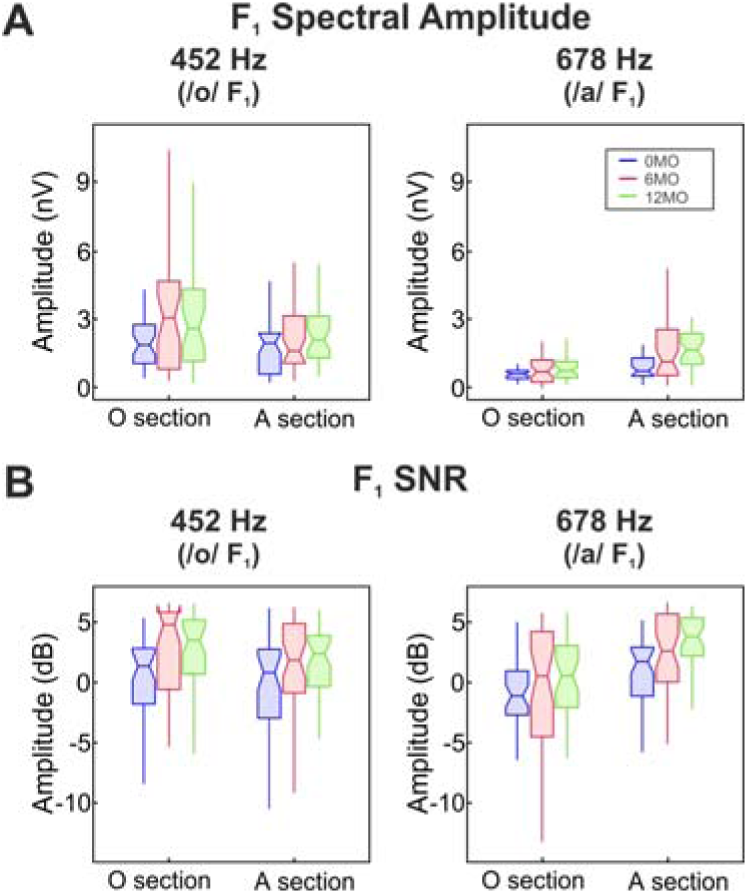
Neural encoding of the first formant corresponding to the two vowel steady sections measured in (**A**) spectral amplitude and (**B**) SNR. Data are plotted from infants at birth (blue), six months (red) and twelve months of age (green), and for /o/ F_1_ (452 Hz, left) and /a/ F_1_ (678 Hz, right) encoding at the different stages of development.

Significant differences in SNR were not observed for the stimulus section (*F*(1,40) = 2.70, *p* = .108, ηp2 = .06). A main effect of age (*F*_(2,80)_ = 8.10, *p* = .001, ηp2 = .17) was obtained with the same pattern as that observed for spectral amplitude. Larger SNRs were found at both six months (*M* = 1.99 ± .57; *p* = .009) and twelve months of age (*M* = 2.10 ± .41; *p* = .001) in comparison to birth (*M* = -.13 ± .53); with no significant variation from six to twelve months of age (*p* = 1). Significant differences were not found for the age per stimulus section interaction (*F*_(2,80)_ = .19, *p* = .825, ηp2 = .005).

#### Neural encoding of the /a/ vowel F_1_

In order to estimate the encoding of the /a/ vowel F_1_ (678 Hz), spectral amplitudes and SNRs were evaluated for the /a/ vowel section. Significant differences were found as a function of age for spectral amplitude (X^2^(2) = 6.78, *p* = .034), with larger values at six months (*Mdn* = .0012; *z* = -2.74, *p* = .006) and twelve months of age (*Mdn* = .0017; *z* =-4.88, *p* < .001) in comparison to values obtained at birth (*Mdn* = .0008). Remarkably, spectral amplitudes obtained at twelve months were also statistically larger than the those at six months (*z* =-2.15, *p* = .032). Similar results were observed for SNR (X^2^(2) = 7.37, *p* = .025), where infants presented larger SNRs at the age of six months (*Mdn* = 2.61; *z* =-2.12, *p* = .034) and at twelve months of age (*Mdn* = 3.81; *z* =-2.84, *p* = .004) compared to birth (*Mdn* = 1.73). No differences in SNR values were observed between six and twelve months of age (*z* = -1.07, *p* = .29). Wilcoxon signed-rank test assessed for the extended six-months old sample (i.e., fifty-four infants) also indicated larger values at six months of age than at birth in the two parameters assessed (spectral amplitude, *z* = -2.67, *p* = .003; SNR, *z* = - 2.31, *p* = .021).

Two-way rmANOVA tests were hence conducted to examine neural response specificity to Stimulus Section (/o/ and /a/) as a function of Age (0, 6 and 12 months) on the spectral amplitude and its SNR at 678 Hz, corresponding to the /a/ F_1_ (see Fig. 4). Spectral amplitude yielded a main effect of stimulus section (*F*_(1,40)_ = 36.36, *p* < .001, ηp2 = .48), with higher values for the /a/ section (*M* = .002 ± <.001) in comparison to the /o/ section (*M* = .001 ± <.001). A main effect of age was also revealed (*F*_(2,80)_ = 7.92, *p* = .001, ηp2 = .17), with significantly larger values at both six months (*M* = .0013 ± <.001; *p* = .010) and twelve months of age (*M* = .0015 ± <.001; *p* = .001) in comparison to the ones obtained at the moment of birth (*M* = .0008 ± <.001); but no significant changes appeared from six to twelve months age (*p* = .89). The age per stimulus section interaction was significant (*F*_(1.74,69.50)_ = 5.99, *p* = .006, ε = .83). *Post-hoc* analysis revealed higher spectral amplitudes at 678 Hz during the /a/ vs. the /o/ vowel sections at the three stages of development (birth, *z* = -3.40, *p* = .001; six-months, *z* = -3.34, *p* = .001; twelve-months, *z* = -4.32, *p* < .001).

SNR results yielded a main effect of stimulus section (*F*_(1,40)_ = 38.90, *p* < .001, ηp2 = .49), indicating higher values at the /a/ section (*M* = 1.93 ± .31) in comparison to the /o/ section (*M* = -.39 ± .30). A main effect of age (*F*_(2,80)_ = 3.83, *p* = .026, ηp2 = .09), with significantly larger values at twelve months of age (*M* = 1.68 ± .42) than at birth (*M* = - .18 ± .43; *p* = .019). No significant age per stimulus section interaction was found (*F*_(1.72,69.06)_ = .80, *p* = .44, ε = .86).

## 4. Discussion

This study describes the longitudinal trajectory of speech-sound neural encoding mechanisms required for language acquisition across the first year of life, as examined through the electrophysiological recordings of the frequency-following response (FFR) elicited by the /oa/ syllable at birth, six and twelve months of age. While no age-related changes were observed in the encoding of voice pitch, a significant enhancement was depicted across the first six-month postnatal period in neural transmission times and neural encoding of the stimulus temporal fine structure. Results contribute to knowledge from previous studies on the developmental trajectory of speech-sound neural encoding mechanisms (Anderson et al., 2015; Arenillas-Alcón et al., 2021, Ribas-Prats et al. 2019, 2023b), by specifically filling the gap with a longitudinal sample spanning the first year of life. Results unveil an early neural maturation in the neural encoding of the speech temporal fine structure and point to a sensitive developmental window in the emergence of core neural mechanisms required for speech acquisition that occurs within the first six months after birth. This neural maturation may underlie the co-occurring critical behavioral language milestones (i.e., acquisition of phonetic categories).

Language acquisition relies on an accurate development of the auditory brain, which is already functional to process sounds at the beginning of the third trimester of pregnancy (Hepper & Shahidullah, 1994; Moore & Linthicum, 2007; Querleu et al., 1988; Ruben, 1995). Around the 27^th^ gestational week, the first traces of myelin can be observed in both the cochlear nerve (Moore & Lithicum, 2001) and the brainstem auditory pathway (Moore, et al., 1995), paralleling the first fetuses’ behavioral and electrophysiological outcomes to auditory stimulation (Draganova et al., 2018; Hepper & Shahidullah, 1994; Schneider et al., 2001). At birth, the cochlea has reached its adult size and is fully functional (Lavigne-Rebillard & Dan Bagger-Sjöbäck, 1992; Moore & Linthicum, 2007), but the auditory brain is not completely mature yet. The shortened neural transmission delay observed in our results at the age of six months aligns with prior literature showing acceleration of auditory neural responses as a function of development (Amorim et al., 2009; Anderson et al., 2015; Madrid et al., 2021; Ribas-Prats et al., 2023b; Sharma et al., 2016). This decrease in neural lag can be attributed to age-related increasing myelination in the auditory white matter tracts along the brainstem, midbrain (Moore et al., 1995), and primary auditory cortex (Su et al., 2008) that occurs during this early period of development.

The perception of speech sound F_0_ and F_1_ plays a crucial role in language acquisition, as these acoustic features constitute key cues that facilitate the learning of a native language (Moon & Hong, 2014). Indeed, voice pitch perception is defined by sound F_0_ (Oxenham, 2012) and provides phonological, syntactic and semantic cues needed for detecting prosodic variation and thus distinguishing word-boundaries from a continuous speech (Nakatani & Schaffer, 1978; Quené, 1993; Rietveld, 1980), as well as for speaker identification (Mary & Yegnanarayana, 2008). Early sensory experience in utero has been demonstrated to be a prerequisite for auditory learning and neural plasticity during the perinatal period (Draganova et al., 2018; Webb et al., 2015). Once they are born, neonates can track the sound envelope, irrespective of the language they have been exposed to during pregnancy (Ortiz Barajas et al., 2021). Our results support previous findings describing an intrauterine perception of low-frequency speech cues (Hepper & Shahidullah, 1994; Voegtline et al., 2013) and an adult-like voice pitch encoding at birth (Anderson et al., 2015; Arenillas-Alcón et al., 2021; Jeng et al., 2011), as no age-related changes were observed here in neither spectral amplitude and SNR values at the stimulus F_0_ peak.

Some discrepancies emerge in literature in relation to the developmental pattern of speech F_0_ neural encoding during the first year of life. While most of the studies report a more robust neural encoding of pitch as a function of age, this pattern does not constantly reach statistical significance across the literature. For instance, Jeng and colleagues (2010) found pitch encoding improvement on a single infant tested at different time-points through the first ten months of age (i.e., 1, 3, 5, 7 and 10 months). Similarly, Van Dyke et al. (2017) described stronger F_0_ neural encoding when comparing a group of older infants (7-12 months old) with a group of younger infants (2-7 months old). Ribas-Prats and colleagues (2023b) also observed age-related improvements from the first postnatal month to six months of age in a longitudinal sample of healthy-term neonates. Yet, in a cross-sectional study performed by Anderson et al. (2015), a similar but not significant trend was found in F_0_ neural encoding for a sample of infants aged from 3 to 10 months. Similarly, our results show a linear pattern of stronger pitch encoding through age during the first postnatal year, although this increase did not reach statistical significance. These discrepancies may arise from the inconsistency on the stimuli frequency components employed across various studies or the individual linguistic environment to which individuals from different study samples are exposed, as suggested by the linguistic experience model (see Jeng et al., 2011; Kuhl et al., 1992). Notably, Jeng and colleagues (2011) compared two samples of neonates exposed to different languages during pregnancy (i.e., English and Chinese) with two matched samples of adults of the same native languages. While Chinese adults showed larger pitch strength values compared to their matched neonates, pitch strength values were comparable between the American neonatal and adult samples. These results seem to highlight a distinctive relevance of pitch encoding in tonal versus non-tonal languages.

Stimulus F_1_ is closely linked to the discrimination of vowel sounds (Kiefte et al., 2010, 2013; Nenadić et al., 2020) and phoneme recognition (Diehl and Lindblom, 2004). Auditory postnatal experience is essential for infants to encode high-frequency components of speech sounds. During pregnancy, the maternal womb acts as a low-pass filter and limits auditory stimulation as it attenuates frequencies above 500 Hz (Gerhardt & Abrahms, 1996, 2000; Hepper & Shahidullah, 1994; Parga et al., 2018), which impedes neonates from hearing high-frequency components before birth. The higher spectral amplitude and SNR values observed at the /a/ vowel F_1_ peak (i.e., 678 Hz) as a function of age support a non-mature neural encoding at birth of these frequency components above circa 500 Hz. Similarly, the ability to track the formant structure of speech seems to be not fully developed at birth, but postnatally experience-dependent, as supported by the higher SNR at both vowels’ F_1_ by the age of six months. These results align with the spectrally ascendant developmental pattern of the auditory system described by Graven and Browne (2008), stating that low-frequency sounds are tuned first in the cochlea, and highlighting the period from 25 gestational weeks to six months of age as the most critical in the neurosensory development of the auditory system. Moreover, the absence of further differences in SNR values at both vowels F_1_ between six and twelve months of age highlights a special relevance of the first six-months postnatal period on the maturation of the temporal fine structure encoding of speech. These results support previous cross-sectional FFR studies reporting an enhancement of F_1_ neural encoding as a function of age (Anderson et al., 2015; Van Dyke et al., 2017), as well as the longitudinal findings by Ribas-Prats et al. (2023b), but further extend these latter findings on the longitudinal trajectory on neural F_1_ encoding through early development to the age of twelve months.

This language-specific attunement to frequencies across the auditory pathway is essential for early language acquisition, as it relies on infants’ ability to apprehend the phonological structure corresponding to a given language (Best et al., 2016; Cutler, 2008). This experience will indeed contribute and facilitate the appropriate identification of native language phonemes by the age of six months (Best et al., 2016; Cheour et al., 1998; Kuhl et al., 1992). Moreover, a perceptual re-organization by the age of six months has been previously proposed, suggesting an attentional shift from syllabic units at birth to phonemic units at six months, cues that are more relevant for word and grammar learning (Nallet & Gervain, 2021; Ortiz Barajas et al., 2021). A rich extrauterine auditory environment is key in the improvement of phonemic categories perception in early language acquisition. During this postnatal period, there is a notable increase in social interactions that play a vital role on infant development, such as in the initial coordination of gestures, vocalizations and facial expressions in interaction with others (Kuhl, 2004, 2010). The co-occurring temporal fine structure encoding bootstrap by the early age of six months revealed in our results suggest an outstanding relevance of this novel and rich extrauterine environment on acoustic and language acquisition.

Future studies are needed to replicate this pattern of development during the first year of life in healthy-term infants. Our results uncover the first six-months of life as a key period in neural speech encoding development. Thus, it is essential to include earlier developmental stages in future longitudinal studies to fully examine this early period of development. Several language-related disabilities such as dyslexia (Banai et al., 2009), learning related disorders (King et al., 2002) or autism (Russo et al., 2008) have been associated with alterations in the spectro-temporal encoding of complex sounds. Also, clinical conditions that occur during the gestational period such as fetal alcohol syndrome (Wyper & Rasmussen, 2011) or fetal growth restriction (Partanen et al., 2018; Ribas-Prats et al., 2022) have an impact on cognitive outcomes, being language one of the major areas affected. Therefore, research on early maturation of neural speech encoding related to language abnormalities in infancy is the next crucial step to comprehend key differences that underlie an inadequate or delayed language acquisition.

Early interventions aimed at improving speech encoding in language affected conditions during the first months of life have not been explored yet. However, the positive effect of musical experience and training in speech encoding has been previously documented for both prenatal (Arenillas-Alcón et al., 2023) and postnatal periods (Wong et al., 2007), suggesting its potential as a promising intervention tool worth exploring. The clinical potential of the electrophysiological tool used in this study has been previously discussed along the literature (Gorina-Careta et al., 2022; Kraus & White-Schwoch, 2015a; Ribas-Prats et al., 2019), suggesting the FFR as a potential biomarker of early language acquisition. Using the FFR as a screening test to early detect speech encoding abnormalities could open the possibility to further design and implement preventive protocols for language-related impairments. The present study provides normative FFR values for the first year of life (i.e., at birth and at six and twelve months of age) and it may thus serve as a reference for future studies on speech-sound neural encoding development.

## 5. Conclusion

The present longitudinal study describes the outstanding maturation of the temporal fine structure neural encoding mechanisms during the very early stages of development. The findings highlight the crucial role of the first six postnatal months in shaping the neural mechanisms that support the encoding of speech sounds, and hence are of major relevance for speech perception and language acquisition. Specifically, our findings unveiled an enhancement in the neural encoding of the formant structure throughout the first six postnatal months, without further maturation up to the first year of life. This reveals a critical maturational period for the neural machinery underlying the ability to discriminate the subtle variations that define phonemes, promoting the formation of phonetic categories. Notably, no significant changes in the neural encoding of voice pitch were observed across this developmental period, which supports a mature voice pitch encoding already at birth. These findings contribute to our understanding of early neural speech encoding and underscore the significance of investigating neural correlates of early speech processing disabilities. Further research in this field can provide valuable guidance for addressing language-related abnormalities and promoting healthy language development in infants.

## Acknowledgements

The authors want to express their gratitude to all the families who generously embraced this project for their selfless participation and commitment to the study.

## References

Aiken, S. J., & Picton, T. W. (2008). Envelope and spectral frequency-following responses to vowel sounds. Hearing Research, 245(1–2), 35–47. 10.1016/j.heares.2008.08.004

Alarcos Llorach, E. (1965). Fonología española [Spanish Phonology]. Gredos.

American Academy of Pediatrics, Joint Committee on Infant Hearing (2019). Year 2019 Position Statement: Principles and Guidelines for Early Hearing Detection and Intervention Programs. The Journal of Early Hearing Detection and Intervention, 4(2), 1–44. https://digitalcommons.usu.edu/jehdi/vol4/iss2/1

Amorim, R. B., Agostinho-Pesse, R. S., & Alvarenga, K. D. F. (2009). The maturational process of the auditory system in the first year of life characterized by brainstem auditory evoked potentials. Journal of Applied Oral Science, 17, 57–62. 10.1590/S1678-77572009000700010

Anderson, S., Parbery-Clark, A., White-Schwoch, T., & Kraus, N. (2015). Development of subcortical speech representation in human infants. The Journal of the Acoustical Society of America, 137(6), 3346–3355. 10.1121/1.4921032

Arenillas-Alcón, S., Costa-Faidella, J., Ribas-Prats, T., Gómez-Roig, M. D., & Escera, C. (2021). Neural encoding of voice pitch and formant structure at birth as revealed by frequency-following responses. Scientific reports, 11(1), 1–16. 10.1038/s41598-021-85799-x

Arenillas-Alcón, S., RibasLPrats, T., Puertollano, M., MondéjarLSegovia, A., GómezLRoig, M. D., CostaLFaidella, J., & Escera, C. (2023). Prenatal daily musical exposure is associated with enhanced neural representation of speech fundamental frequency: Evidence from neonatal frequencyLfollowing responses. Developmental Science, 00, e13362. 10.1111/desc.13362

Banai, K., Hornickel, J., Skoe, E., Nicol, T., Zecker, S., & Kraus, N. (2009). Reading and subcortical auditory function. Cerebral cortex, 19(11), 2699–2707. 10.1093/cercor/bhp024

Banai, K., Nicol, T., Zecker, S. G., & Kraus, N. (2005). Brainstem timing: implications for cortical processing and literacy. Journal of Neuroscience, 25(43), 9850–9857. 10.1523/JNEUROSCI.2373-05.2005

Basu, M., Krishnan, A., & WeberLFox, C. (2010). Brainstem correlates of temporal auditory processing in children with specific language impairment. Developmental science, 13(1), 77–91. 10.1111/j.1467-7687.2009.00849.x

Best, C. T., Goldstein, L. M., Nam, H., & Tyler, M. D. (2016). Articulating what infants attune to in native speech. Ecological Psychology, 28(4), 216–261. 10.1080/10407413.2016.1230372

Byers-Heinlein, K., Burns, T. C., & Werker, J. F. (2010). The roots of bilingualism in newborns. Psychological science, 21(3), 343–348. 10.1177/0956797609360758

Cheour, M., Ceponiene, R., Lehtokoski, A., Luuk, A., Allik, J., Alho, K., & Näätänen, R. (1998). Development of language-specific phoneme representations in the infant brain. Nature neuroscience, 1(5), 351–353. 10.1038/1561

Coffey, E. B., Nicol, T., White-Schwoch, T., Chandrasekaran, B., Krizman, J., Skoe, E., Zatorre, R. J., & Kraus, N. (2019). Evolving perspectives on the sources of the frequency-following response. Nature communications, 10(1), 5036. 10.1038/s41467-019-13003-w

Cooley, J. W., & Tukey, J. W. (1965). An algorithm for the machine calculation of complex Fourier series. Mathematics of computation, 19(90), 297–301. 10.2307/2003354

Corp., I. SPSS 25.0. (Corp., I, Chicago). https://www.ibm.com/

Cunningham, J., Nicol, T., Zecker, S. G., Bradlow, A., & Kraus, N. (2001). Neurobiologic responses to speech in noise in children with learning problems: deficits and strategies for improvement. Clinical Neurophysiology, 112(5), 758–767. 10.1016/s1388-2457(01)00465-5

Cutler, A. (2008). The 34th Sir Frederick Bartlett Lecture: The abstract representations in speech processing. Quarterly journal of experimental psychology, 61(11), 1601–1619. 10.1080/13803390802218542

DeCasper, A. J., & Fifer, W. P. (1980). Of human bonding: Newborns prefer their mothers’ voices. Science, 208(4448), 1174–1176. 10.1126/science.7375928

Diehl, R.L., Lindblom, B. (2004). Explaining the Structure of Feature and Phoneme Inventories: The Role of Auditory Distinctiveness. In: Speech Processing in the Auditory System. Springer Handbook of Auditory Research, vol 18. Springer, New York, NY. 10.1007/0-387-21575-1_3

Draganova, R., Schollbach, A., Schleger, F., Braendle, J., Brucker, S., Abele, H., Kagan, K. O., Wallwiener, D., Fritsche, A., Eswaran, H., & Preissl, H. (2018). Fetal auditory evoked responses to onset of amplitude modulated sounds. A fetal magnetoencephalography (fMEG) study. Hearing Research, 363, 70–77. 10.1016/j.heares.2018.03.005

Feldman, H. M. (2019). How young children learn language and speech. Pediatrics in review, 40(8), 398–411. 10.1542/pir.2017-0325

Ferreira, L., Skarzynski, P. H., Skarzynska, M. B., Sanfins, M. D., & Biaggio, E. P. V. (2021). Effect of Auditory Maturation on the Encoding of a Speech Syllable in the First Days of Life. Brain Sciences, 11(7), 844. 10.3390/brainsci11070844

Figueras, F., & Gratacós, E. (2014). Update on the diagnosis and classification of fetal growth restriction and proposal of a stage-based management protocol. Fetal diagnosis and therapy, 36(2), 86–98. 10.1159/000357592

Font-Alaminos, M., Cornella, M., Costa-Faidella, J., Hervás, A., Leung, S., Rueda, I., & Escera, C. (2020). Increased subcortical neural responses to repeating auditory stimulation in children with autism spectrum disorder. Biological psychology, 149, 107807. 10.1016/j.biopsycho.2019.107807

Gerhardt, K. J., & Abrams, R. M. (1996). Fetal hearing: characterization of the stimulus and response. Seminars in perinatology, 20(1), 11–20. 10.1016/s0146-0005(96)80053-x

Gerhardt, K. J., & Abrams, R. M. (2000). Fetal exposures to sound and vibroacoustic stimulation. Journal of Perinatology, 20(1), S21–S30. 10.1038/sj.jp.7200446

Gorina-Careta, N., Kurkela, J. L., Hämäläinen, J., Astikainen, P., & Escera, C. (2021). Neural generators of the frequency-following response elicited to stimuli of low and high frequency: A magnetoencephalographic (MEG) study. Neuroimage, 231, 117866. 10.1016/j.neuroimage.2021.117866

Gorina-Careta, N., Ribas-Prats, T., Arenillas-Alcón, S., Puertollano, M., Gómez-Roig, M. D., & Escera, C. (2022). Neonatal frequency-following responses: A methodological framework for clinical applications. Seminars in Hearing, 43(03), 162–176. 10.1055/s-0042-1756162

Granier-Deferre, C., Bassereau, S., Ribeiro, A., Jacquet, A. Y., & DeCasper, A. J. (2011). A melodic contour repeatedly experienced by human near-term fetuses elicits a profound cardiac reaction one month after birth. PLoS One, 6(2), e17304. 10.1371/journal.pone.0017304

Graven, S. N., & Browne, J. V. (2008). Auditory development in the fetus and infant. Newborn and infant nursing reviews, 8(4), 187–193. 10.1053/j.nainr.2008.10.010

Hepper, P. G., Scott, D., & Shahidullah, S. (1993). Newborn and fetal response to maternal voice. Journal of Reproductive and Infant Psychology, 11(3), 147–153. 10.1080/02646839308403210

Hepper, P. G., & Shahidullah, B. S. (1994). The development of fetal hearing. Fetal and Maternal Medicine Review, 6(3), 167–179. 10.1017/S0965539500001108

Hervé, E., Mento, G., Desnous, B., & François, C. (2022). Challenges and new perspectives of developmental cognitive EEG studies. NeuroImage, 260, 119508. 10.1016/j.neuroimage.2022.119508

Hornickel, J., & Kraus, N. (2013). Unstable representation of sound: a biological marker of dyslexia. Journal of Neuroscience, 33(8), 3500–3504. 10.1523/JNEUROSCI.4205-12.2013

Hualde, J. I. (2013). Los sonidos del español: Spanish language edition [The sounds of Spanish: Spanish language edition]. Cambridge University Press.

Jeng, F. C., Hu, J., Dickman, B., Montgomery-Reagan, K., Tong, M., Wu, G., & Lin, C. D. (2011). Cross-linguistic comparison of frequency-following responses to voice pitch in American and Chinese neonates and adults. Ear and hearing, 32(6), 699–707. 10.1097/AUD.0b013e31821cc0df

Jeng, F. C., Lin, C. D., & Wang, T. C. (2016). Subcortical neural representation to Mandarin pitch contours in American and Chinese newborns. The Journal of the Acoustical Society of America, 139(6), EL190. 10.1121/1.4953998

Jeng, F. C., Schnabel, E. A., Dickman, B. M., Hu, J., Li, X., Lin, C. D., & Chung, H. K. (2010). Early maturation of frequency-following responses to voice pitch in infants with normal hearing. Perceptual and motor skills, 111(3), 765–784. 10.2466/10.22.24.PMS.111.6.765-784

Jusczyk, P.W., Luce, P.A., & Charles-Luce, J. (1994). Infants’ sensitivity to phonotactic patterns in the native language. Journal of Memory and Language, 33, 630–645. 10.1006/jmla.1994.1030

Kiefte, M., Enright, T., & Marshall, L. (2010). The role of formant amplitude in the perception of/i/and/u. The Journal of the Acoustical Society of America, 127(4), 2611–2621. 10.1121/1.3353124

Kiefte, M., Nearey, T. M., Assmann, P. F., Ball, M. J., & Gibbon, F. E. (2013). Vowel perception in normal speakers. Handbook of vowels and vowel disorders, 2, 160.

King, C., Warrier, C. M., Hayes, E., & Kraus, N. (2002). Deficits in auditory brainstem pathway encoding of speech sounds in children with learning problems. Neuroscience letters, 319(2), 111–115. 10.1016/S0304-3940(01)02556-3

Krizman, J., & Kraus, N. (2019). Analyzing the FFR: A tutorial for decoding the richness of auditory function. Hearing research, 382, 107779. 10.1016/j.heares.2019.107779

Kuhl, P. K. (2004). Early language acquisition: cracking the speech code. Nature reviews neuroscience, 5(11), 831–843. 10.1038/nrn1533

Kuhl, P. K. (2010). Brain mechanisms in early language acquisition. Neuron, 67(5), 713–727. 10.1016/j.neuron.2010.08.038

Kuhl, P. K., Stevens, E., Hayashi, A., Deguchi, T., Kiritani, S., & Iverson, P. (2006). Infants show a facilitation effect for native language phonetic perception between 6 and 12 months. Developmental science, 9(2), F13–F21. 10.1111/j.1467-7687.2006.00468.x

Kuhl, P. K., Tsao, F. M., & Liu, H. M. (2003). Foreign-language experience in infancy: Effects of short-term exposure and social interaction on phonetic learning. Proceedings of the National Academy of Sciences, 100(15), 9096–9101. 10.1073/pnas.1532872100

Kuhl, P. K., Williams, K. A., Lacerda, F., Stevens, K. N., & Lindblom, B. (1992). Linguistic experience alters phonetic perception in infants by 6 months of age. Science, 255(5044), 606–608. 10.1126/science.1736364

Kujala, T., Partanen, E., Virtala, P., & Winkler, I. (2023). Prerequisites of language acquisition in the newborn brain. Trends in Neurosciences. 10.1016/j.tins.2023.05.011

Lavigne-Rebillard, M., & Bagger-Sjöbäck, D. (1992). Development of the human stria vascularis. Hearing research, 64(1), 39–51. 10.1016/0378-5955(92)90166-k

Madrid, A. M., Walker, K. A., Smith, S. B., Hood, L. J., & Prieve, B. A. (2021). Relationships between click auditory brainstem response and speech frequency following response with development in infants born preterm. Hearing Research, 407, 108277. 10.1016/j.heares.2021.108277

Martí i Roca, J. M. (1986). Paràmetres acústics per a la síntesi de consonants fricatives catalanes [Acoustic parameters for Catalan fricative consonants’ synthesis]. Estudios de fonética experimental, 151–193.

Mary, L., & Yegnanarayana, B. (2008). Extraction and representation of prosodic features for language and speaker recognition. Speech communication, 50(10), 782–796. 10.1016/j.specom.2008.04.010

Maye, J., Werker, J. F., & Gerken, L. (2002). Infant sensitivity to distributional information can affect phonetic discrimination. Cognition, 82(3), B101–B111. 10.1016/s0010-0277(01)00157-3

Mehler, J., Jusczyk, P., Lambertz, G., Halsted, N., Bertoncini, J., & Amiel-Tison, C. (1988). A precursor of language acquisition in young infants. Cognition, 29(2), 143–178. 10.1016/0010-0277(88)90035-2

Moon, I. J., & Hong, S. H. (2014). What is temporal fine structure and why is it important?. Korean journal of audiology, 18(1), 1. 10.7874/kja.2014.18.1.1

Moore, J. K., & Linthicum Jr, F. H. (2001). Myelination of the human auditory nerve: different time courses for Schwann cell and glial myelin. *Annals of Otology*, Rhinology & Laryngology, 110(7), 655–661. 10.1177/000348940111000711

Moore, J. K., & Linthicum Jr, F. H. (2007). The human auditory system: a timeline of development. International journal of audiology, 46(9), 460–478. 10.1080/14992020701383019

Moore, J. K., Perazzo, L. M., & Braun, A. (1995). Time course of axonal myelination in the human brainstem auditory pathway. Hearing Research, 91(1–2), 208–209. 10.1016/0378-5955(95)00218-9

Nakatani, L. H., & Schaffer, J. A. (1978). Hearing “words” without words: Prosodic cues for word perception. The Journal of the Acoustical Society of America, 63(1), 234–245. 10.1121/1.381719

Nallet, C., & Gervain, J. (2021). Neurodevelopmental preparedness for language in the neonatal brain. Annual Review of Developmental Psychology, 3, 41–58. 10.1146/annurev-devpsych-050620-025732

Nazzi, T., Bertoncini, J., & Mehler, J. (1998). Language discrimination by newborns: toward an understanding of the role of rhythm. Journal of Experimental Psychology: Human perception and performance, 24(3), 756. 10.1037//0096-1523.24.3.756

Nenadić, F., Coulter, P., Nearey, T. M., & Kiefte, M. (2020). Perception of vowels with missing formant peaks. The Journal of the Acoustical Society of America, 148(4), 1911–1921. 10.1121/10.0002110

Ortiz Barajas, M. C., Guevara, R., & Gervain, J. (2021). The origins and development of speech envelope tracking during the first months of life. Developmental cognitive neuroscience, 48, 100915. 10.1016/j.dcn.2021.100915

Otto-Meyer, S., Krizman, J., White-Schwoch, T., & Kraus, N. (2018). Children with autism spectrum disorder have unstable neural responses to sound. Experimental Brain Research, 236(3), 733–743. 10.1007/s00221-017-5164-4

Oxenham, A. J. (2012). Pitch perception. Journal of Neuroscience, 32(39), 13335–13338. 10.1523/JNEUROSCI.3815-12.2012

Parga, J. J., Daland, R., Kesavan, K., Macey, P. M., Zeltzer, L., & Harper, R. M. (2018). A description of externally recorded womb sounds in human subjects during gestation. PloS one, 13(5), e0197045. 10.1371/journal.pone.0197045

Partanen, L., Korkalainen, N., Mäkikallio, K., Olsén, P., LaukkanenLNevala, P., & Yliherva, A. (2018). Foetal growth restriction is associated with poor reading and spelling skills at eight years to 10 years of age. Acta Paediatrica, 107(1), 79–85. 10.1111/apa.14005

Quené, H. (1993). Segment durations and accent as cues to word segmentation in Dutch. The Journal of the Acoustical Society of America, 94(4), 2027–2035. 10.1121/1.407504

Querleu, D., Renard, X., Versyp, F., Paris-Delrue, L., & Crèpin, G. (1988). Fetal hearing. European Journal of Obstetrics & Gynecology and Reproductive Biology, 28(3), 191–212. 10.1016/0028-2243(88)90030-5

Ribas-Prats, T., Almeida, L., Costa-Faidella, J., Plana, M., Corral, M. J., Gómez-Roig, M. D., & Escera, C. (2019). The frequency-following response (FFR) to speech stimuli: A normative dataset in healthy newborns. Hearing Research, 371, 28–39. 10.1016/j.heares.2018.11.001

Ribas-Prats, T., ArenillasLAlcón, S., LipLSosa, D. L., CostaLFaidella, J., Mazarico, E., GómezLRoig, M. D., & Escera, C. (2022). Deficient neural encoding of speech sounds in term neonates born after fetal growth restriction. Developmental Science, 25(3), e13189. 10.1111/desc.13189

Ribas-Prats, T., Arenillas-Alcón, S., Pérez-Cruz, M., Costa-Faidella, J., Gómez-Roig, M. D., & Escera, C. (2023a). Speech-encoding deficits in neonates born large-for-gestational age as revealed with the envelope frequency-following response. Ear and hearing 44(4), 829–841. 10.1097/AUD.0000000000001330

Ribas-Prats, T., Cordero, G., Lip-Sosa, D. L., Arenillas-Alcón, S., Costa-Faidella, J., Gómez-Roig, M. D., & Escera, C. (2023b). Developmental Trajectory of the Frequency-Following Response During the First 6 Months of Life. *Journal of Speech*, Language, and Hearing Research, 66(12), 4785–4800. 10.1044/2023_JSLHR-23-00104

Rietveld, A. C. M. (1980). Word boundaries in the French language. Language and Speech, 23(3), 289–296. 10.1177/002383098002300306

Rivera Gaxiola, M., Silva Pereyra, J., & Kuhl, P. K. (2005). Brain potentials to native and non native speech contrasts in 7 and 11 month old American infants. Developmental science, 8(2), 162–172. 10.1111/j.1467-7687.2005.00403.x

Ruben, R. J. (1995). The ontogeny of human hearing. International journal of pediatric otorhinolaryngology, 32, S199–S204. 10.1016/0165-5876(94)01159-U

Russo, N. M., Skoe, E., Trommer, B., Nicol, T., Zecker, S., Bradlow, A., & Kraus, N. (2008). Deficient brainstem encoding of pitch in children with autism spectrum disorders. Clinical Neurophysiology: official journal of the International Federation of Clinical Neurophysiology, 119(8), 1720–1731. 10.1016/j.clinph.2008.01.108

Schneider, U., Schleussner, E., Haueisen, J., Nowak, H., & Seewald, H. J. (2001). Signal analysis of auditory evoked cortical fields in fetal magnetoencephalography. Brain topography, 14, 69–80. 10.1023/A:1012519923583

Sharma, M., Bist, S. S., & Kumar, S. (2016). Age-related maturation of wave V latency of auditory brainstem response in children. Journal of Audiology & Otology, 20(2), 97. 10.7874/jao.2016.20.2.97

Stuart, A., Yang, E. Y., & Green, W. B. (1994). Neonatal auditory brainstem response thresholds to air-and bone-conducted clicks: 0 to 96 hours postpartum. Journal of the American Academy of Audiology, 5(3), 163–172.

Su, P., Kuan, C. C., Kaga, K., Sano, M., & Mima, K. (2008). Myelination progression in language-correlated regions in brain of normal children determined by quantitative MRI assessment. International Journal of Pediatric Otorhinolaryngology, 72(12), 1751–1763. 10.1016/j.ijporl.2008.05.017

The MathWorks Inc. (2019). MATLAB version: 9.7 (R2019b), Natick, Massachusetts: The MathWorks Inc. https://www.mathworks.com

Tsao, F. M., Liu, H. M., & Kuhl, P. K. (2004). Speech perception in infancy predicts language development in the second year of life: A longitudinal study. Child development, 75(4), 1067–1084. 10.1111/j.1467-8624.2004.00726.x

Tsao, F. M., Liu, H. M., & Kuhl, P. K. (2006). Perception of native and non-native affricate-fricative contrasts: Cross-language tests on adults and infants. The Journal of the Acoustical Society of America, 120(4), 2285–2294. 10.1121/1.2338290

Van Dyke, K. B., Lieberman, R., Presacco, A., & Anderson, S. (2017). Development of phase locking and frequency representation in the infant frequency-following response. *Journal of Speech*, Language, and Hearing Research, 60(9), 2740–2751. 10.1044/2017_JSLHR-H-16-0263

Voegtline, K. M., Costigan, K. A., Pater, H. A., & DiPietro, J. A. (2013). Near-term fetal response to maternal spoken voice. Infant Behavior and Development, 36(4), 526–533. 10.1016/j.infbeh.2013.05.002

Webb, A. R., Heller, H. T., Benson, C. B., & Lahav, A. (2015). Mother’s voice and heartbeat sounds elicit auditory plasticity in the human brain before full gestation. Proceedings of the National Academy of Sciences, 112(10), 3152–3157. 10.1073/pnas.1414924112

Werker, J. F., Gilbert, J. H., Humphrey, K., & Tees, R. C. (1981). Developmental aspects of cross-language speech perception. Child development, 349-355. 10.2307/1129249

Werker, J. F., & Tees, R. C. (1984). Cross-language speech perception: Evidence for perceptual reorganization during the first year of life. Infant behavior and development, 7(1), 49–63. 10.1016/S0163-6383(84)80022-3

White-Schwoch, T., Davies, E. C., Thompson, E. C., Carr, K. W., Nicol, T., Bradlow, A. R., & Kraus, N. (2015b). Auditory-neurophysiological responses to speech during early childhood: Effects of background noise. Hearing research, 328, 34–47. 10.1016/j.heares.2015.06.009

White-Schwoch, T., Woodruff Carr, K., Thompson, E. C., Anderson, S., Nicol, T., Bradlow, A. R., Zecker, S. G., & Kraus, N. (2015a). Auditory processing in noise: A preschool biomarker for literacy. PLoS biology, 13(7), e1002196. 10.1371/journal.pbio.1002196

Wong, P. C., Skoe, E., Russo, N. M., Dees, T., & Kraus, N. (2007). Musical experience shapes human brainstem encoding of linguistic pitch patterns. Nature neuroscience, 10(4), 420–422. 10.1038/nn1872

Wyper, K. R., & Rasmussen, C. R. (2011). Language impairments in children with fetal alcohol spectrum disorder. Journal of Population Therapeutics and Clinical Pharmacology, 18(2). Retrieved from https://www.jptcp.com/index.php/jptcp/article/view/485

